# Bacterial division FtsZ forms liquid condensates with nucleoid-associated Z-ring inhibitor SlmA

**DOI:** 10.1101/264192

**Authors:** Begoña Monterroso, Silvia Zorrilla, Marta Sobrinos-Sanguino, Miguel A. Robles-Ramos, Marina López-Álvarez, Christine D. Keating, Germán Rivas

**Affiliations:** Centro de Investigaciones Biológicas, Consejo Superior de Investigaciones Científicas (CSIC) 28040, Madrid, Spain; Department of Chemistry, Pennsylvania State University, University Park, Pennsylvania 16802, USA.

**Author notes:** These authors contributed equally to this work (Begoña Monterroso, Silvia Zorrilla). For correspondence (BM), (SZ), (GR).

## Abstract

Macromolecular condensation resulting from biologically regulated liquid-liquid phase transitions is emerging as a mechanism to organize the intracellular space in eukaryotic systems, with broad implications in cell physiology and pathology. Here we show that FtsZ, central element of the division ring in most bacteria, forms condensates when in complex with SlmA, the protein preventing septal ring assembly nearby the chromosome in *E. coli*. The formation of condensates is promoted by crowding and enhanced by sequence-specific binding of SlmA to DNA. These structures are dynamic and FtsZ within them remains active for GTP-triggered fiber formation. Their location is sensitive to compartmentalization and to the presence of a membrane boundary in microfluidics-based cell mimetic systems, likely affecting their reactivity. We propose that reversible condensation may play a role in the modulation of FtsZ assembly and/or location by SlmA and, hence, in the regulation of ring stability, constituting a singular example of a prokaryotic nucleoprotein complex exhibiting this kind of phase transition.

## Introduction

Recent studies evidence the relevance of the transient condensation of proteins and other molecules into structures behaving as liquid droplets in the spatiotemporal regulation of biological processes [1]. The emergence of these membraneless compartments can be described as a liquid demixing process generating phase boundaries to temporarily confine specific functional entities [2]. The selective accumulation of biomolecules in these dynamic compartments may deeply influence their reactivity, enhancing and accelerating their mutual recognition while disfavoring interactions with excluded elements [3-5]. Crowding due to the overall high concentration of macromolecules in the cells clearly provides a non-specific driving force favoring phase transitions and condensation [6-10], another contributing factor being, it seems, multivalency [1, 3, 11, 12]. Indeed, most biomolecular condensates analyzed consist of various molecules containing multiple homo or heteroassociation elements, like nucleic acids, usually RNA, and proteins harboring various domains of interaction [1]. The presence of unstructured regions in proteins also seems to promote condensation and a number of intrinsically disordered proteins have been found to form liquid droplets on their own under crowding conditions *in vitro* and *in vivo* that evolve towards solid aggregates [9, 10, 13-15]. Specific features shared by these condensates, like the ability to exchange molecules with the surroundings, their local and reversible generation in response to changes in component interactions and/or concentrations, and their evolving physical properties [1] make them particularly suitable for the fine tuning of molecular localization and reactivity.

Phase separation has been shown to occur in a variety of signaling and other systems in eukaryotes (see, for instance, works by Woodruff [8], Courchaine [16] and Hennig [17]). In prokaryotes, however, the membraneless organelles described drop to a few examples that, with the exception of the bacterial nucleoid hypothesized to be a liquid phase [18-20], are structurally different from the eukaryotic condensates. Among them, carboxysomes and Pdu microcompartments [21-23], resembling a virus shell encircling a number of enzymes that participate in linked reactions, are able to exchange metabolites with the surrounding cytoplasm through the pores of this capsid [24]. It remains to be determined if prokaryotic proteins may transiently condensate into dynamic structures like those described for a growing number of eukaryotic systems and, if so, the factors driving this condensation and their possible functional implications.

Bacterial division is a key process for cell survival in which many multivalent interactions of proteins, nucleic acids and lipids occur, hence being an attractive system to investigate the effect of phase transitions and crowding in prokaryotes. Division is achieved through the assembly of a dynamic ring at midcell that constricts the membrane, giving rise to two daughter cells of similar size and identical genomic content [25]. The scaffold of this ring is built on filaments of FtsZ, a GTPase that self-associates in the presence of GTP, able to interact with other division proteins, among them modulators of the localization of the ring [26, 27]. FtsZ assembly results in the formation of dynamic single stranded filaments that come apart upon GTP depletion due to hydrolysis [26, 28]. In the presence of homogeneous or heterogeneous crowders, whether inert polymers, DNA or proteins, mimicking the agglomeration in the cytoplasm, excluded volume effects and other non-specific interactions enhance the tendency of the protein to assemble into filaments [29] that can interact laterally to form fibers [30, 31]. Reconstruction of FtsZ in liquid-liquid phase separation (LLPS) systems mimicking microenvironments and compartmentalization, in bulk or encapsulated in microdroplets, shows that the protein distributes unevenly among phases and/or interfaces and this distribution can be reversibly modulated by its association state [32, 33].

Bacteria have developed different mechanisms to control the formation of the division ring in space and time [34]. One of these mechanisms is nucleoid occlusion, mediated in *E. coli* by the DNA binding protein SlmA, which precludes Z-ring formation in the surroundings of the chromosome protecting it from scission upon septum formation [27]. SlmA, when in complex with its specifically recognized palindromic DNA sequences (SBSs), counteracts the assembly of FtsZ into filaments [35, 36] accelerating their disassembly [37]. Contrary to other antagonists that sequester FtsZ subunits decreasing the GTPase activity [38-40] and blocking polymerization all over the cell, SlmA modulation has a spatial dimension. Thus, it occurs selectively nearby all chromosomal macrodomains but the SBS-free Ter one [35, 41] without changing the GTPase activity of FtsZ single stranded filaments [37, 42].

Despite the mechanistic insights provided by recent research on SlmA and its role in division [43], there is no information available on the impact of this protein on the reactivity and organization of FtsZ species in crowded media. Here we have analyzed how SlmA and its complexes with a short double stranded DNA containing the consensus sequence specifically recognized by the protein (SBS) modulate the behavior of FtsZ in crowding conditions and in LLPS systems mimicking cellular microenvironments. The FtsZ·SlmA division complexes were also reconstructed in microfluidics-based microdroplets, containing an LLPS system and stabilized by a lipid mixture matching the composition in the *E. coli* inner membrane (EcL), as cell-like environments displaying a membrane boundary and compartmentalization. Our results show that FtsZ complexes with SlmA form structures consistent with crowding driven liquid droplets that evolve towards fibers upon GTP addition and reassemble with GTP exhaustion. This constitutes a unique example of a nucleoprotein system in prokaryotes displaying a transient arrangement into dynamic liquid droplets, which might be important for its function, as recently described for some protein-nucleic acid systems in eukaryotes.

## Results

### FtsZ forms dynamic condensates upon interaction with SlmA under crowding conditions

The effect of the nucleoid occlusion protein SlmA on the behavior of FtsZ in crowding conditions was initially probed using dextran 500 as a crowding agent. Confocal images of a sample containing FtsZ labelled with Alexa 647 (FtsZ-Alexa 647), SlmA and a fluorescein labelled 24 bp double stranded oligonucleotide containing the consensus sequence targeted by SlmA (SBS-Fl) in dextran showed abundant round structures in which the two dyes colocalized (**Fig. 1AB**), reminiscent of liquid droplets (1-6 μm diameter). Such structures were not found to be formed by either FtsZ in the absence of the inhibitory complex or by FtsZ·SlmA·SBS in the absence of crowders (**Fig. 1C**, **Supplementary Fig. 1**). Numerous round FtsZ·SlmA·SBS condensates were also observed in the presence of high concentrations (50-150 g/L) of other typically used crowders like Ficoll 70 and PEG 8 (**Fig. 1DE**). Turbidity measurements further confirmed the formation of these structures in solutions containing the crowding agents but not in their absence (**Fig. 1F**). When only SlmA was added to FtsZ in crowding conditions, condensates in which the two labelled proteins colocalized were also found, although in lower amounts (**Supplementary Fig. 2**). Therefore crowding, by volume exclusion and/or other unspecific effects, provides a driving force for the formation of droplet-like structures of FtsZ in the presence of SlmA and this type of arrangement is strongly favored by specific binding to the SBS oligonucleotide.

**Fig. 1.**
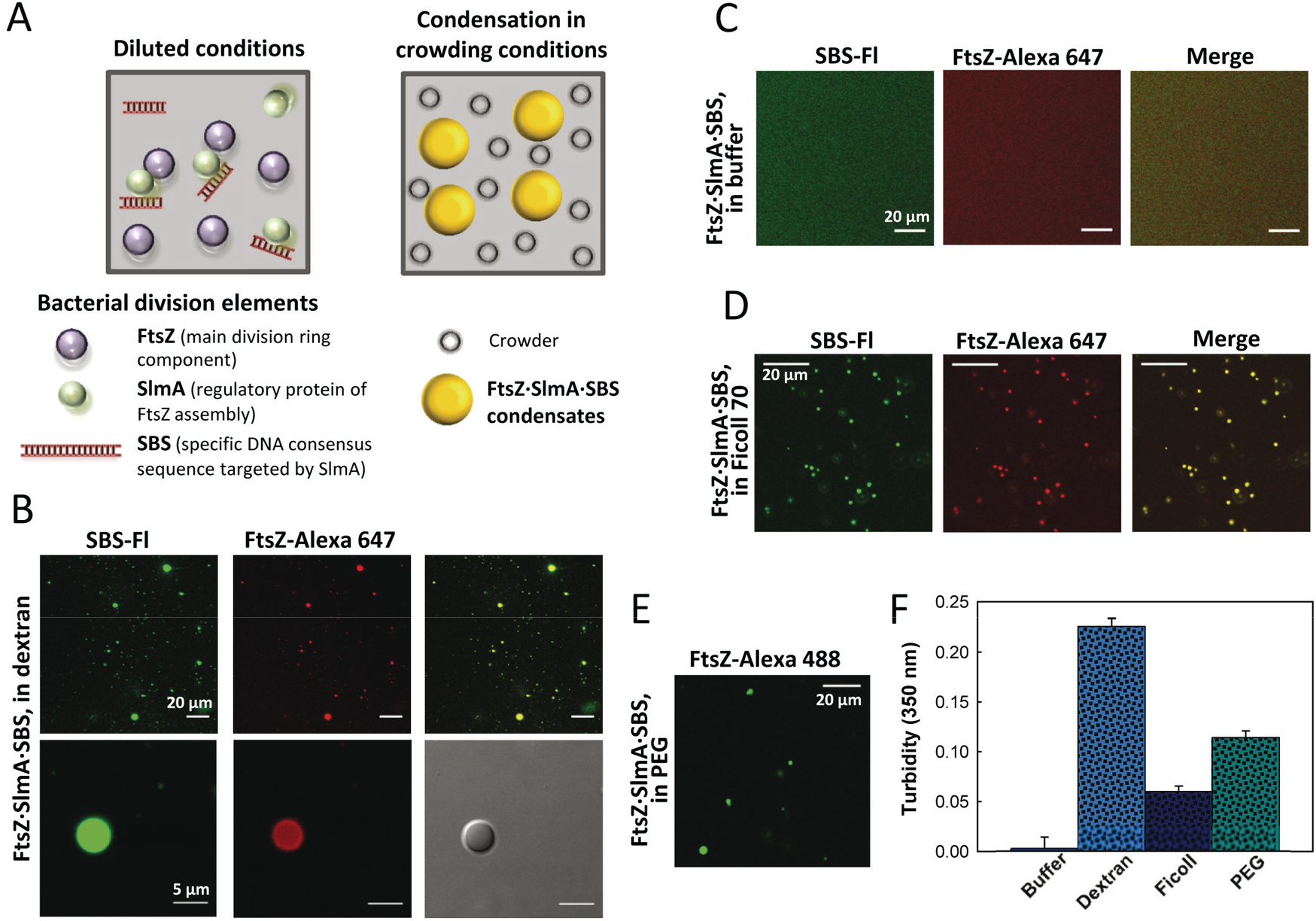
Formation of condensates of FtsZ and SlmA·SBS in crowding conditions. (**A**) Scheme of the *E. coli* division elements involved in the formation of condensates. (**B**) Representative confocal images of the FtsZ·SlmA·SBS (25 μM FtsZ) condensates in 80 g/L dextran. Images at higher magnification are shown below. (**C**) Absence of condensates in dilute solution. (**D**) FtsZ·SlmA·SBS condensates in 150 g/L Ficoll and (**E**) in 50 g/L PEG (25 μM FtsZ). (**F**) Turbidity of FtsZ·SlmA·SBS in buffer, in 150 g/L dextran 500 or Ficoll 70 and in 50 g/L PEG 8. The concentrations of FtsZ, SlmA and SBS in all panels were 12, 5 and 1 μM, unless when otherwise stated.

To further characterize these round assemblies we studied whether they were dynamic, a characteristic feature of liquid droplets, for which we performed experiments of FtsZ capture similar to those recently reported for other protein systems forming condensates [8]. Images show the final state and temporal evolution of nucleoprotein FtsZ·SlmA·SBS complexes containing FtsZ labelled with Alexa 647 in 150 g/L dextran to which FtsZ-Alexa 488 was subsequently added (**Fig. 2A**, **Supplementary Movie 1**). Colocalization of the two dyes showed the recruitment of new FtsZ into the preformed FtsZ·SlmA·SBS round condensates indicating that, being the overall arrangement maintained, FtsZ within these structures was dynamic. The condensates of FtsZ·SlmA·SBS remained dynamic and able to incorporate newly added FtsZ after more than 3 hours within similar time scales as the freshly prepared samples (**Supplementary Fig. 3**). Diffusion of FtsZ into FtsZ·SlmA·SBS complexes was also found in other crowding solutions like Ficoll and PEG, showing a similar behavior and comparable times of protein uptake among them (**Fig. 2BC**) and with those observed in dextran. Analogous FtsZ diffusion experiments performed with samples containing FtsZ·SlmA condensates (*i.e.* in the absence of the SBS oligonucleotide) showed colocalization of the green and red labelled FtsZ within seconds, despite of the fewer droplets observed (**Supplementary Fig. 4**, **Supplementary Movie 2**). Thus, the round structures formed by FtsZ in the presence of SlmA with or without the specific SBS oligonucleotide sequence in crowding conditions behave as permeable dynamic condensates that exchange protein with the surroundings, as expected for liquid droplets.

**Fig. 2.**
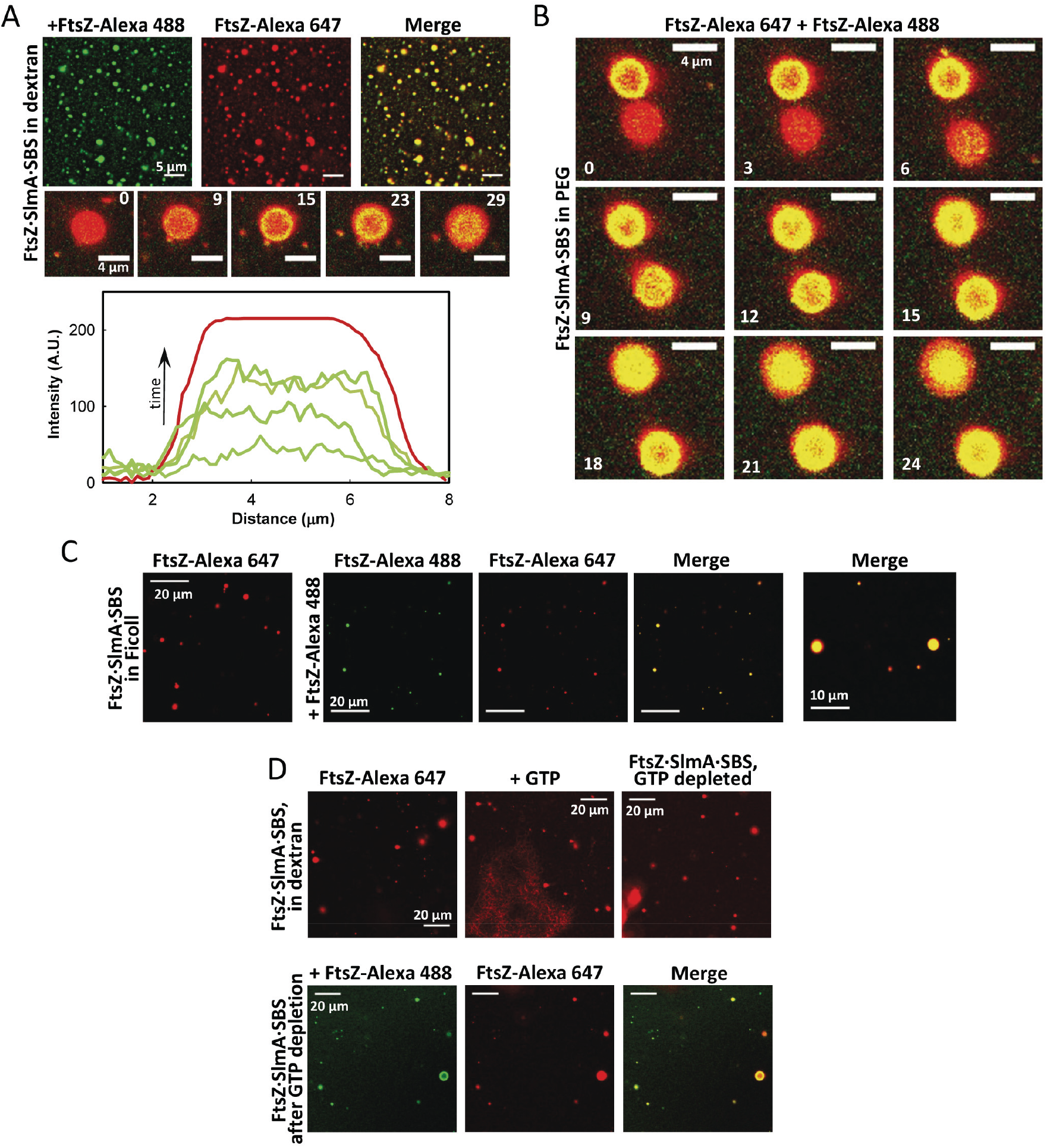
Dynamism of FtsZ·SlmA·SBS condensates. Representative confocal images showing (**A**) Final state after addition of FtsZ-Alexa 488 to FtsZ·SlmA·SBS complexes (FtsZ labelled with Alexa 647) in 150 g/L dextran. Below, images showing the stepwise diffusion of FtsZ-Alexa 488 into the FtsZ·SlmA·SBS droplets containing FtsZ-Alexa 647 at the indicated times in seconds (time zero, beginning of visualization for that particular droplet) and corresponding intensity profiles at selected times in the green channel. The profile in the red channel is shown as a reference and varies slightly among images. (**B**) Stepwise diffusion of FtsZ-Alexa 488 into FtsZ·SlmA·SBS condensates (FtsZ labelled with Alexa 647) at the indicated times in seconds (time zero, beginning of visualization for those particular droplets) in 50 g/L PEG. (**C**) Initial (far left panel) and final states after diffusion of FtsZ-Alexa 488 into the condensates of FtsZ·SlmA·SBS containing FtsZ-Alexa 647 in 150 g/L Ficoll. An image of the final state (merge) with higher magnification is included on the right. (**D**) Assembly of FtsZ within FtsZ·SlmA·SBS condensates into fibers upon GTP addition (0.7 mM) and condensates formed after FtsZ fibers disassembly in 150 g/L dextran. Below, final state after addition of FtsZ-Alexa 488 on condensates formed by FtsZ·SlmA·SBS (FtsZ labelled with Alexa 647) after FtsZ fibers disassembly due to GTP depletion, in 150 g/L dextran.

Next we asked if FtsZ within the condensates remained active for assembly into fibers. Addition of GTP on previously formed droplets of FtsZ·SlmA·SBS nucleoprotein complexes in dextran readily induced the formation of FtsZ fibers and, upon disassembly of the fibers, the two proteins and the oligonucleotide incorporated back into liquid droplets (**Fig. 2D**). The condensates obtained after GTP depletion maintained the dynamic nature observed before triggering of FtsZ assembly, as stated by time lapse imaging of protein incorporation showing the newly added FtsZ rapidly colocalizing with the condensates labelled with a spectrally different dye (**Fig. 2D**). These experiments reinforce the idea that the condensates are dynamic entities in which FtsZ retains its GTP dependent self-association and hydrolysis properties.

### Compartmentalization affects the distribution and localization of the condensates formed by FtsZ and SlmA

To determine the effect that microenvironments as those found in the cell might have on the condensates formed by FtsZ in the presence of SlmA, bound to its target SBS sequence, we chose a mixture widely characterized among the LLPS systems, PEG 8/dextran 500, in which unassembled FtsZ distributes asymmetrically, partitioning preferentially into the dextran rich phase [32]. Confocal images of an emulsion of PEG/dextran containing SlmA·SBS and FtsZ showed abundant condensates, of size similar to those formed in the homogeneous crowders (∼2 μm diameter), in which the two proteins and the specific SBS oligonucleotide colocalize (**Fig. 3AB**, **Supplementary Fig. 5A**). The condensates distributed mostly within dextran, as shown by the lack of colocalization with labelled PEG (**Supplementary Fig. 5B**). These structures were not found with FtsZ alone [32], fewer condensates were observed when only SlmA was added to FtsZ (**Supplementary Fig. 6AB**) and they were virtually nonexistent when FtsZ was in the presence of free SBS (**Supplementary Fig. 7A**, top row). Scarce condensates were occasionally found in samples with SlmA or SlmA·SBS in the PEG/dextran LLPS system (**Supplementary Fig. 7BC**) indicating that, although SlmA itself could somehow maintain the ability to form droplets on its own, it is in the presence of FtsZ when these structures abound in the solutions. Images were acquired with different combinations of dyes and the FtsZ·SlmA condensates, with or without SBS oligonucleotide, were observed irrespectively of the dye and of which element was labelled (**Supplementary Figs. 5** and **6**).

**Fig. 3.**
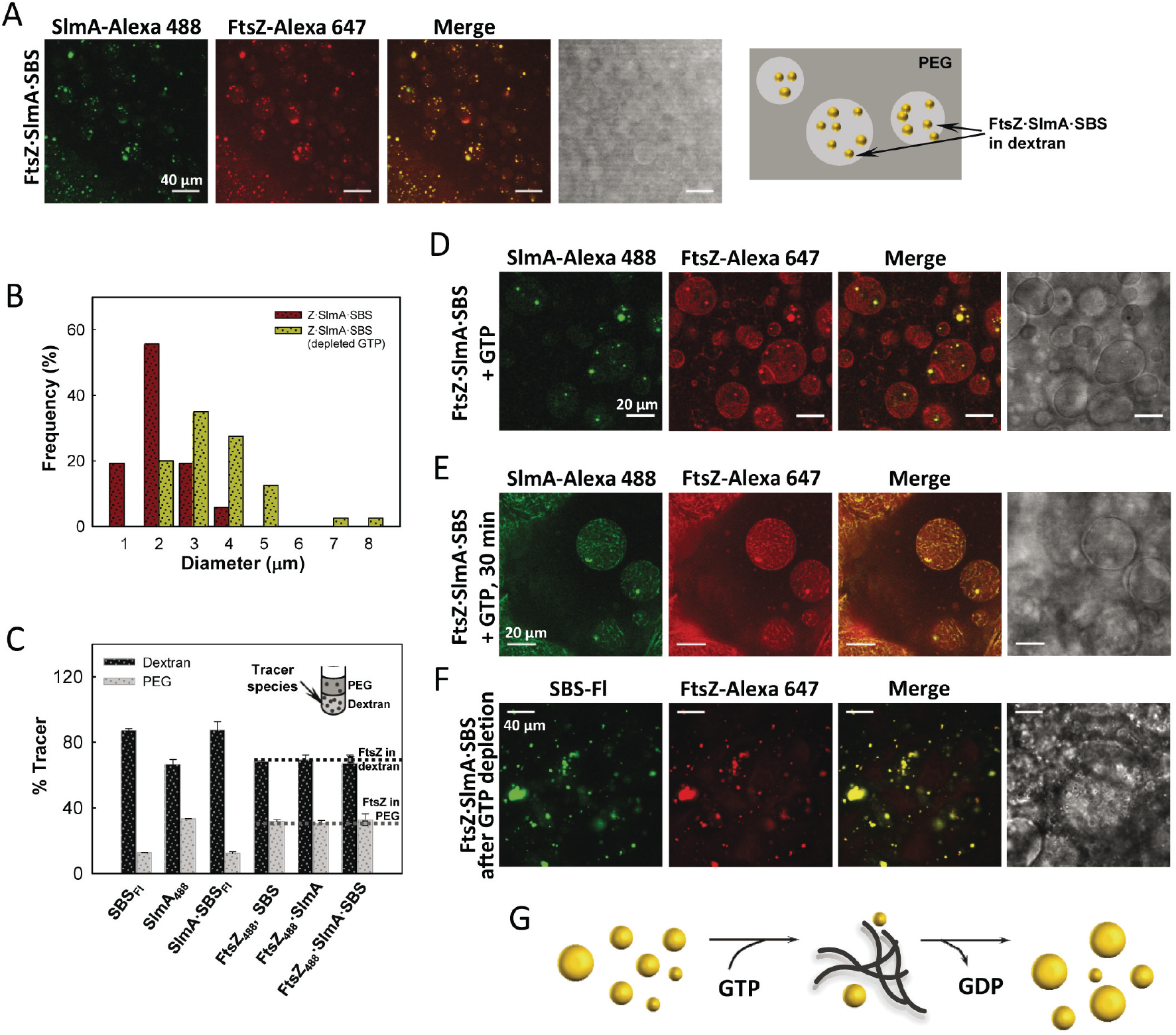
Formation of FtsZ·SlmA·SBS condensates in the PEG/dextran LLPS system and evolution upon GTP-induced FtsZ assembly into fibers. (**A**) Representative confocal and transmitted images of FtsZ·SlmA·SBS complexes and scheme of their distribution within the LLPS system on the right. (**B**) Distribution of sizes of FtsZ·SlmA·SBS liquid droplets before (*n* = 52) and after (*n* = 40) an FtsZ fiber assembly/disassembly cycle. (**C**) Partition of SlmA, SBS and the complexes with FtsZ within the LLPS mixture as determined by fluorescence, together with an illustration. Horizontal lines depict, for comparison, distribution within these phases of FtsZ alone. (**D**) FtsZ·SlmA·SBS complexes in the presence of GTP (1 mM) at time zero and (**E**) 30 min after addition of the nucleotide. (**F**) FtsZ·SlmA·SBS condensates formed after FtsZ fiber disassembly due to GTP (0.33 mM) depletion. (**G**) Scheme of the dynamic process entailing FtsZ·SlmA·SBS condensates and GTP induced fibers. When GTP is added to samples containing FtsZ·SlmA·SBS condensates fibers are formed and the number of condensates decrease. The condensates rearrange upon GTP depletion and fibers disassembly.

Fluorescence measurements of total protein partition showed that the distribution of FtsZ, predominantly (although not exclusively) in the dextran (**Fig. 3C**) was not substantially altered upon interaction with SlmA·SBS or by the presence of free SBS oligonucleotide or SlmA, both with a preference, specially marked in the case of SBS, for dextran (**Fig. 3C**, **Supplementary Fig. 7CD**). The condensates observed in the images accumulated in the dextran phase, disregarding whether they include the specific SBS oligonucleotide or not, likely because of the higher concentration of protein in this phase.

### The specific nucleoprotein complexes of SlmA modulate the arrangement and localization of FtsZ fibers in LLPS systems mimicking compartmentalization

As the role of SlmA in division is the negative modulation of the GTP dependent FtsZ assembly, we analyzed the impact of its condensates with FtsZ on the ability of the latter to assemble into fibers in LLPS systems and on the distribution of the fibers eventually formed. Addition of GTP on the FtsZ·SlmA·SBS complexes in PEG/dextran readily induced FtsZ assembly into filaments decorated with SlmA and SBS oligonucleotide (**Fig. 3DE**, **Supplementary Fig. 8**), apparently smaller than those formed in this LLPS system by FtsZ alone (**Supplementary Fig. 7F**, top row). The fibers distributed mostly within the dextran phase and, although not excluded from the interface, their preference for this location was not as blatant as that of fibers formed solely by FtsZ (**Fig. 3D**, **Supplementary Fig. 7F**), likely due to their smaller size as a result of the action of SlmA·SBS. Monitoring of the evolution of the sample with time showed that it initially contained a significant number of nucleoprotein condensates coexisting with fibers, and as the fibers became more abundant the number and average size of the droplets decreased (**Fig. 3DE**). Depletion of GTP seemed to reverse the process as, upon disassembly, liquid condensates slightly bigger on average than those present before inducing fiber formation were found (∼3.5 μm diameter; **Fig. 3BFG**). Control experiments showed that the presence of SBS oligonucleotide, without SlmA, had no apparent effect on the general aspect and usual location of FtsZ fibers [32], mostly in dextran and at the interface (*cf.* **Supplementary Figs. 7F**, top row, and **7A**, bottom row). In the presence of SlmA, with no SBS, they appeared somewhat thinner and their preference for the interface was reduced (**Supplementary Fig. 7E**), suggesting that SlmA by itself could have an antagonistic effect on FtsZ assembly in crowding conditions, as previously described in diluted solution at high concentrations [35].

We also studied the effect of SlmA·SBS complexes on preformed FtsZ fibers in the binary PEG/dextran system. Addition of the antagonist complex immediately rendered thinner and apparently shorter FtsZ filaments that redistributed within both PEG and dextran phases and the formation, afterwards, of liquid condensates located within the dextran phase (**Supplementary Fig. 7F**). These experiments show that SlmA bound to its specific nucleic acid sequence modify the general arrangement and distribution of FtsZ fibers in LLPS systems mimicking compartmentalization, and further confirmed that the formation of liquid droplets in dynamic equilibrium with fibers, modulated by GTP binding and hydrolysis, is inherent to the FtsZ·SlmA·SBS system (**Fig. 3G**).

### Liquid condensates of FtsZ and SlmA accumulate at lipid surfaces

To determine how the membrane boundary and confinement in the *E. coli* cells may impact the formation of condensates by FtsZ and SlmA we reconstructed the two proteins and the SBS sequence within microenvironments inside microdroplets, picoliter cell mimic systems surrounded by a lipid boundary, using microfluidics based technology. Following an approach we recently optimized [33], we induced the formation of the FtsZ·SlmA·SBS complex in PEG/dextran by simultaneously encapsulating in microdroplets, stabilized by the *E. coli* lipid mixture, FtsZ (with a tracer amount of FtsZ-Alexa 647) in the stream containing the PEG solution and SlmA·SBS (SBS oligonucleotide labelled with fluorescein) in that with the dextran solution (**Fig. 4A**). Round condensates where observed inside the microdroplets, mostly at or nearby the lipid interface, and the specific SBS sequence and FtsZ outside these condensates were mainly located in one of the phases, presumably the dextran, the preference of SBS for this phase being more marked (**Fig. 4B**). We next encapsulated the same solutions including GTP in the stream with dextran, triggering the formation of FtsZ fibers shortly before their actual encapsulation. Around 30 minutes after production, FtsZ was almost completely disassembled and showed basically the distribution observed with the oligomeric form. The arrangement into condensates in which FtsZ and the SBS colocalized was evident shortly after, appearing mainly at and nearby the lipid interface (**Fig. 4C**, **Supplementary Movie 3**) and increasing their number with time. In order to observe the fibers and characterize the distribution of species before nucleotide depletion, we followed the same procedure for the encapsulation but increasing the nucleotide concentration and reducing those of FtsZ and SlmA·SBS, thus increasing the lifetime of the fibers (**Fig. 4D**). Images show the presence, within the dextran phase and at the interface, of FtsZ filaments thinner than in the absence of the inhibitory complex [33] in which SlmA and FtsZ colocalized. The proteins (and the complexed SBS oligonucleotide) were also present at the lipid boundary and, in those areas of the microdroplet showing a higher local concentration of SlmA·SBS, the abundance of filaments decreased. Interestingly enough, the size of the confined liquid droplets, whether formed after depolymerization or not, was smaller (∼1 μm diameter) than those observed under the same conditions in bulk solution (*cf*. **Figs. 3** and **4**). These experiments indicate that the FtsZ·SlmA·SBS system retains the tendency to reversibly form condensates also when encapsulated inside micron-sized containers mimicking the compartmentalization of the cytoplasm. Moreover, these condensates may have an impact on the distribution and localization of the two proteins as they seem to display a marked tendency towards the lipid boundary.

**Fig. 4.**
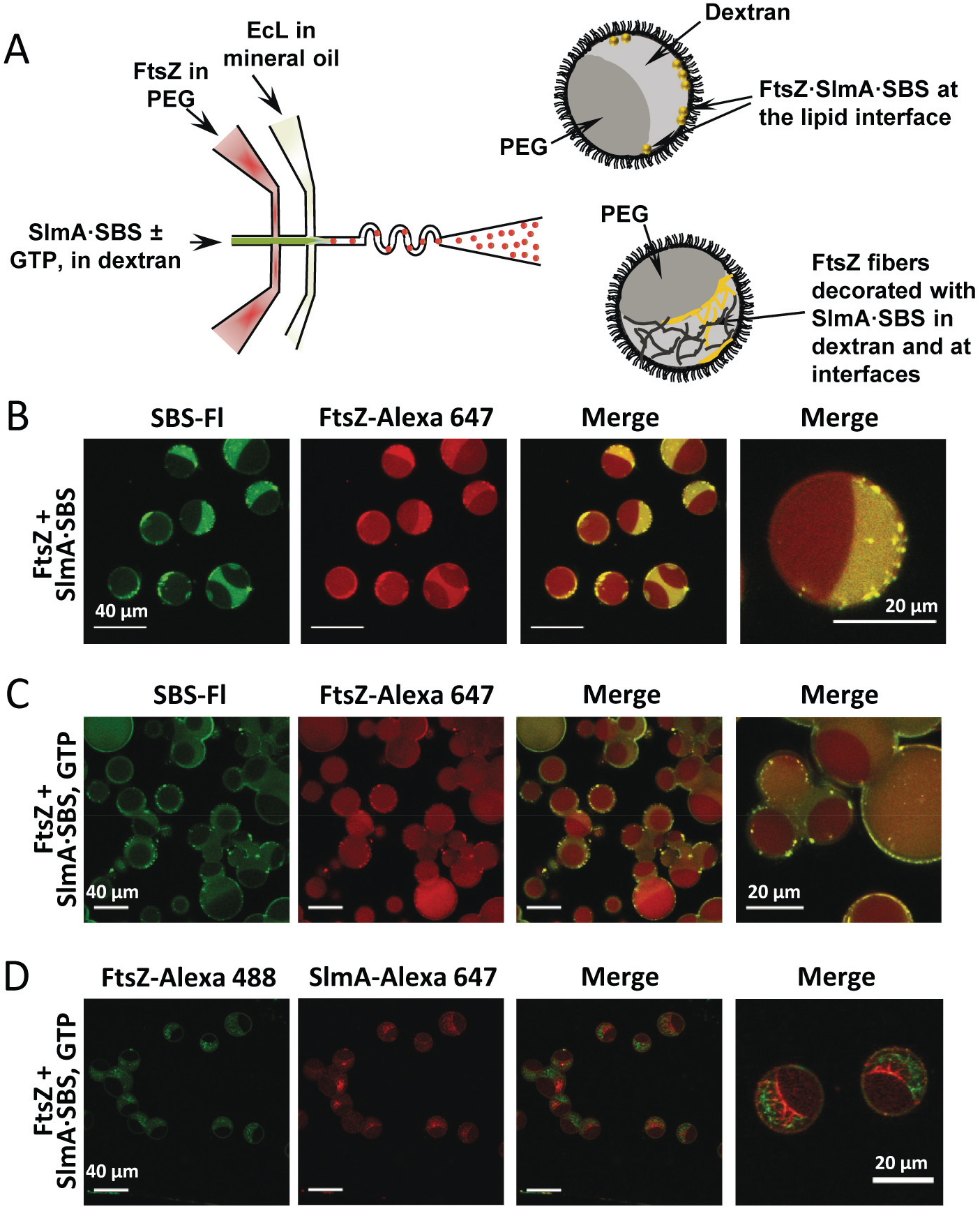
Microfluidic encapsulation of FtsZ·SlmA·SBS in the PEG/dextran LLPS system inside microdroplets stabilized by the *E. coli* lipid mixture. (**A**) Scheme of the encapsulation setup and illustration, on the right, of the distribution of species within the encapsulated LLPS system. (**B-D**) Representative confocal images of the microdroplets without (B) and with GTP (C and D). The concentrations of FtsZ, SlmA and SBS were 12, 5 and 1 μM respectively (B and C) or 6, 3 and 0.5 μM respectively (D). 1 mM (C) or 2 mM GTP (D).

### SlmA driven condensation of FtsZ in PEG/DNA as a model LLPS system closer to an intracellular environment

One LLPS system particularly relevant in the case of the DNA binding protein SlmA, in which we have previously studied the behavior of FtsZ [32], is that consisting of mixtures of PEG and nucleic acid phases. The nucleic acid phase consists of short salmon sperm DNA fragments prepared as previously described [44], allowing to reproduce some of the features of nucleic acid rich compartments in the bacterial cytoplasm as the charged nature. In this PEG/DNA system FtsZ, SlmA and the specific SBS oligonucleotide targeted by the latter also condensed into liquid droplet-like structures and they were especially numerous and substantially bigger (3-4 μm diameter) than those formed in PEG/dextran, distributing preferentially (but not exclusively) in the DNA phase (**Fig. 5A**, **Supplementary Fig. 9A**). Each of the individual components of the complex showed a clear preferential distribution for the DNA, in the case of SBS not as remarkable but still obvious (**Supplementary Fig. 9B**). FtsZ samples in PEG/DNA also contained droplet-like structures when SlmA was solely added, located within the DNA phase and more abundant than those in PEG/dextran (*cf.* **Supplementary Figs. 6A** and **9C**), likely because of the unspecific charge effects of the DNA at high concentration and/or unspecific binding of SlmA to these sequences. The condensates were, however, apparently smaller than when the specific SBS oligonucleotide was also present (∼2 μm). No condensates were found for FtsZ in the absence of SlmA [32]. In the absence of FtsZ, SlmA was able to form liquid droplets when bound to the SBS oligonucleotide, although in a very limited number and, without SBS, SlmA only rarely formed them (**Supplementary Fig. 9B**), suggesting that the simultaneous presence of SlmA, FtsZ and DNA, preferably the specific SBS sequence, is determinant to achieve significant condensation. Similarly than with other crowders, the FtsZ·SlmA·SBS condensates obtained at high concentrations of the unspecific DNA in the LLPS system were dynamic, as they captured FtsZ freshly added to the sample (**Fig. 5B**). These results indicate that the formation of liquid droplets by the division proteins occurs not only in the presence of inert crowding agents but also of negatively charged ones mimicking the high nucleic acid content of the bacterial cytoplasm. Moreover, as in the PEG/dextran system, the locally higher concentrations of the components due to partitioning into one of the phases, in this case the DNA-rich one, would probably further enhance condensation.

**Fig. 5.**
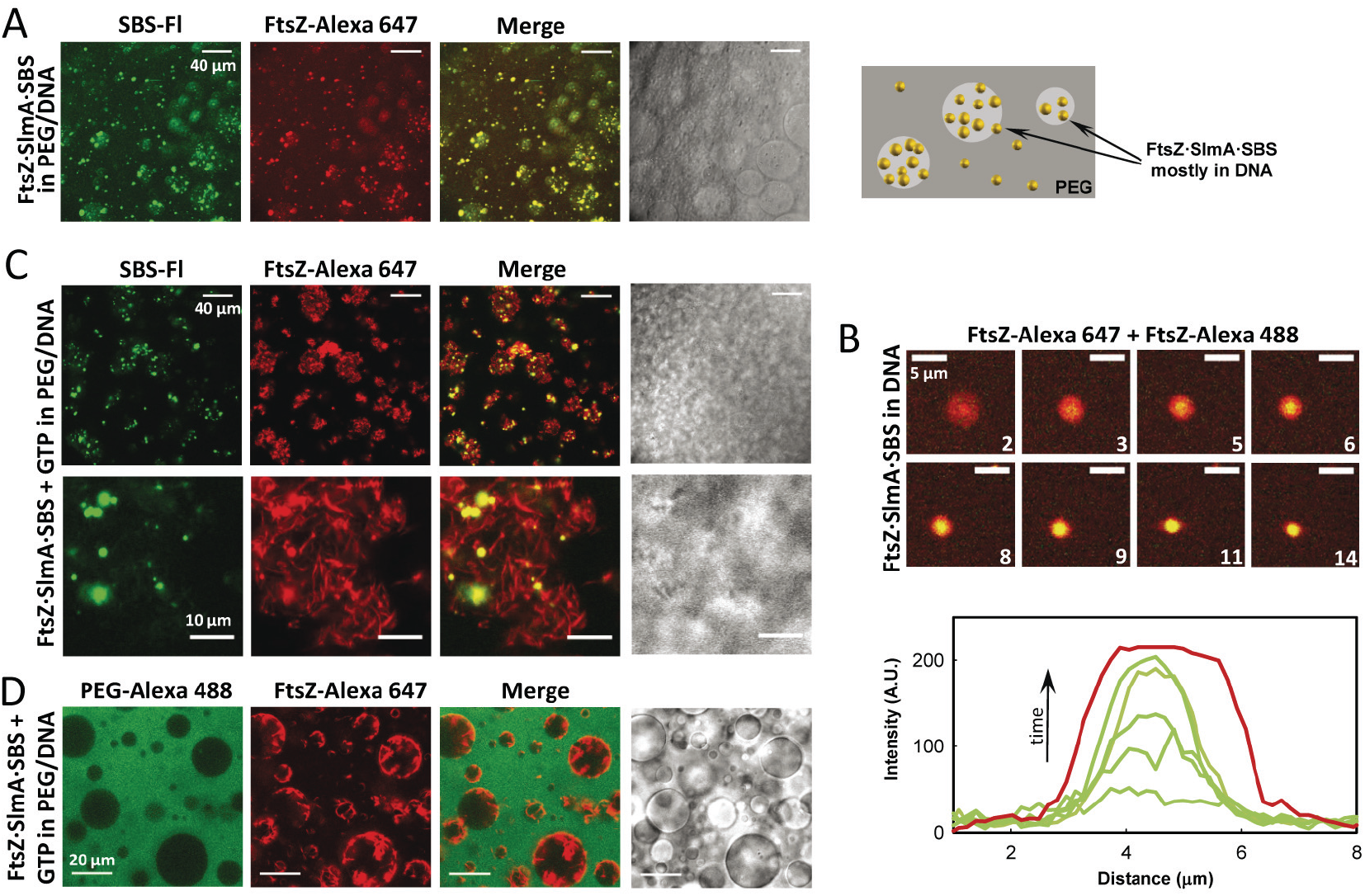
FtsZ·SlmA·SBS condensates and GTP induced fibers in the PEG/DNA LLPS system. (**A**) Representative confocal images of FtsZ·SlmA·SBS complexes and, on the right, schematic illustration of their disposition within the phases. (**B**) Stepwise diffusion of FtsZ-Alexa 488 added on FtsZ·SlmA·SBS condensates (FtsZ labelled with Alexa 647) at the indicated times in seconds (time zero, beginning of visualization for this particular droplet) in 180 g/L DNA, and representative intensity profiles in the green channel at different times below. The profile in the red channel, shown as a reference, varies slightly within the images. (**C**) and (**D**) Changes induced in FtsZ·SlmA·SBS complexes upon GTP addition (2 mM).

We next checked if FtsZ in the condensates was still active for fiber formation in the PEG/DNA LLPS system. With GTP addition, numerous and highly thickened fibers of FtsZ coexisting with large liquid droplets appeared mainly (and randomly) distributed within the DNA phase and at the interface (**Fig. 5CD**). Although the three elements seem to be colocalizing in both types of structures, stronger signal was developed within the condensates (**Fig. 5C**). The presence of SlmA and its specific SBS oligonucleotide did not alter the preferential distribution of the FtsZ fibers in the DNA phase [32]. However, the fibers appeared more evenly distributed within this phase occupying all the available space, including the interface (**Supplementary Movie 4**), likely because of the influence of the inhibitory SlmA complex on their size and arrangement.

The formation of condensates of the division proteins was also analyzed by reconstruction in microfluidics-based microdroplets stabilized by the *E. coli* lipid mixture containing the PEG/DNA LLPS system to reproduce charged and uncharged compartments. Encapsulation of the PEG/DNA LLPS system required slight adaptation of the procedure earlier described [33]. Remarkably, encapsulation in this case was more straightforward than with the PEG/dextran LLPS system, probably because of the lower viscosity of the DNA solution. We started characterizing the behavior of FtsZ, either assembled or not, including the protein in the stream containing PEG, the other aqueous stream containing the unspecific DNA crowder and, when required, GTP. The microdroplets generated were of similar size (∼15-20 μm diameter) as those previously produced with the PEG/dextran mixture and the distribution of both phases among them was homogeneous (**Supplementary Fig. 10**). The partition of the protein in the microdroplets was similar to that previously obtained in the PEG/DNA system encapsulated by simple emulsion [32] with the protein mainly in the DNA phase, in the case of the fibers concentrated in some areas of this phase leaving others practically devoid of them, and some protein also at the lipid surface (**Supplementary Fig. 10**).

Next, to determine the effect of the SlmA·SBS complex on FtsZ we included it in the unspecific DNA stream (**Fig. 6A**). In the absence of GTP, condensates in which FtsZ and the SBS oligonucleotide colocalized were observed and, as with the PEG/dextran LLPS system, they were mainly located at the lipid interface, and occasionally also at the interface between both crowders (**Fig. 6B**, **Supplementary Movie 5**). The protein and SBS out of the condensates remained distributed within the phases, preferentially at the DNA. In the presence of GTP, fibers of FtsZ, decorated by labelled SBS presumably in complex with SlmA, were observed mainly in the DNA, and a large fraction of the protein and of the SBS located at the lipids (**Fig. 6C**). Condensates were still not visible after around 30 minutes, although discontinuities in the colocalization pattern in the membrane could be observed (**Fig. 6C**). With time, however, FtsZ fibers seem to gather and slowly migrate towards the lipid boundary (**Supplementary Movies 6** and **7**), maybe in response to partial structural changes corresponding to initial stages in the formation of the condensates upon GTP depletion.

**Fig. 6.**
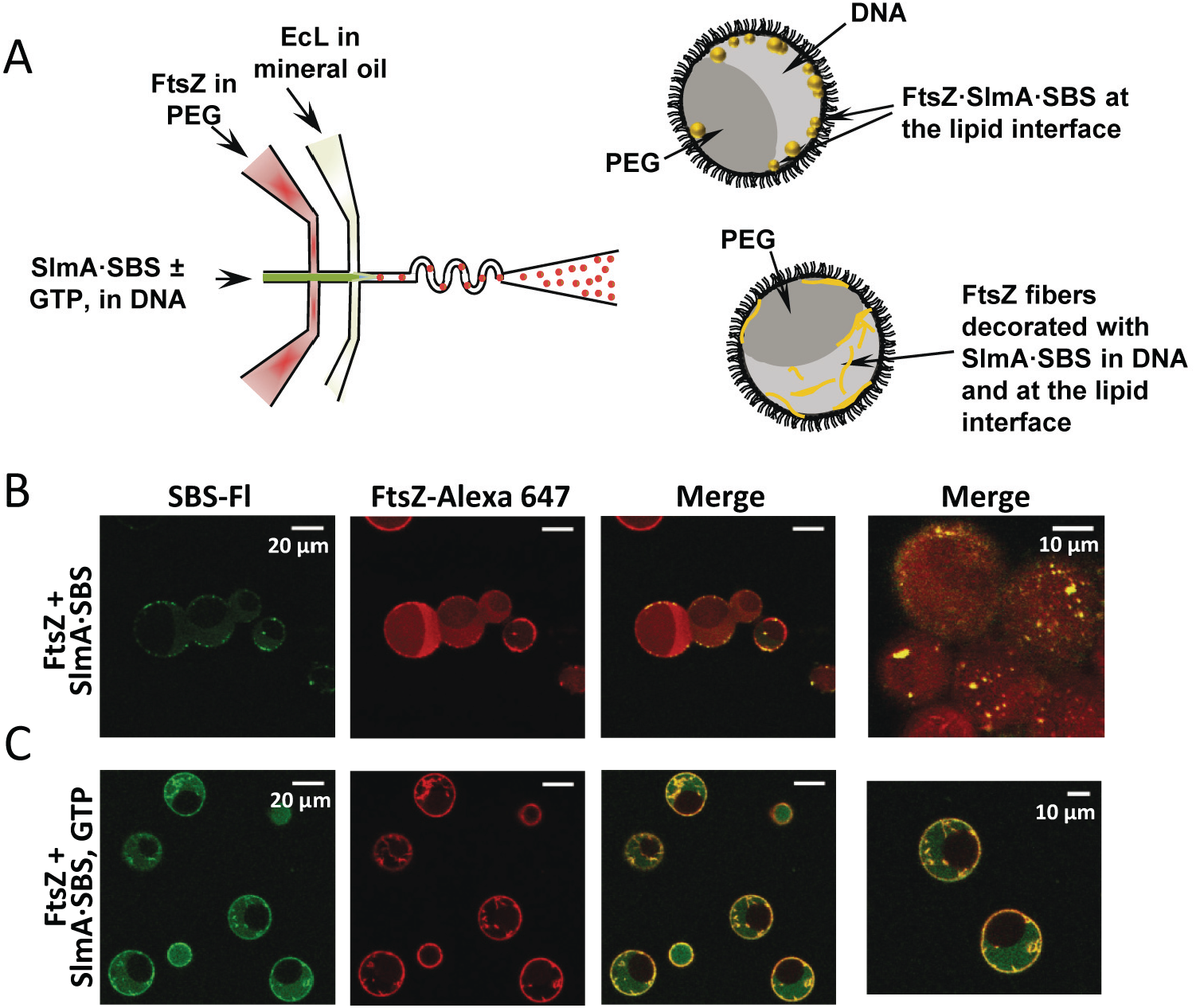
Microfluidic encapsulation of FtsZ·SlmA·SBS in the PEG/DNA LLPS system inside microdroplets and effect of GTP induced FtsZ fiber formation. (**A**) Scheme of the encapsulation procedure followed for the PEG/DNA LLPS system and illustration, on the right, of the distribution of species within the encapsulated LLPS system. (**B**) and (**C**) Representative confocal images of the microdroplets stabilized by the *E. coli* lipid mixture containing the biphasic PEG/DNA mixture and the complex FtsZ·SlmA·SBS without and with 2 mM GTP, respectively. Last image in (B) focuses in the lipid interface to show the high density of condensates. Concentrations were 12 μM FtsZ, 5 μM SlmA, 1 μM SBS.

## Discussion

Here we show that, under crowding conditions and in LLPS systems mimicking microenvironments, including those encapsulated by microfluidics as cell models, FtsZ reversibly forms dynamic condensates when in the presence of SlmA in complex with its specific SBS oligonucleotide sequences, and that these condensates are consistent with crowding-induced liquid droplets. The generation of compartments usually involved in diverse metabolic pathways has been previously described in prokaryotes. Examples of the few structures of this kind so far identified are carboxysomes, Pdu and Eut microcompartments, which are multiprotein complexes encased in a porous protein shell allowing the limited exchange of substrates and reaction products with the surroundings [21-23]. The dynamic condensates of the division proteins observed here are somehow different as they only require a discrete number of components to be assembled and none of the molecules involved seems to provide a capsid encircling the whole structure, as they capture externally added protein. Therefore, rather than similar to the prokaryotic microcompartments so far known, these new structures are reminiscent of the membraneless eukaryotic condensates recently described for intrinsically disordered proteins and multivalent complexes, the assembly of which is promoted by crowding [1, 3, 8-11].

Condensates of FtsZ in the presence of SlmA and, eventually, also of SlmA bound to its specific SBS sequence can be found in crowding conditions. However, the formation of liquid droplets is more obvious when the three elements are present, consistent with the idea that multivalent interactions favor condensation [1, 3, 11]. Multivalency is clearly present in the FtsZ·SlmA·SBS complexes [43], as the two proteins involved contain multiple domains of homo- and heteroassociation, and the nucleic acid sequence anchors, in diluted solution, a dimer of dimers of SlmA [37, 42]. Although FtsZ forms discrete oligomers in the absence of GTP [45], greatly enhanced in size and number under crowding conditions [46], the interaction with SlmA and preferably also with the SBS is required for liquid droplet formation, in line with previous observations that nucleic acids favor this type of structures [1, 9, 13]. Together with multivalency, crowding seems to be the other factor providing a driving force for FtsZ·SlmA·SBS condensation, as observed for other proteins forming liquid droplets in eukaryotes [8, 13], presumably because of the decrease of available volume and/or other non-specific interactions favoring macromolecular association [47]. All of the inert crowders tested here induce condensates but their abundance and size seems to be dramatically enhanced at high concentrations of DNA, particularly when phase separated from PEG. This is likely due to the additional exclusion provoked by electrostatic repulsion between the DNA molecules themselves and with FtsZ units and the specific SBS sequences, which are also negatively charged at the working pH. Large effects arising from high concentrations of DNA on FtsZ assembly and organization have been described before [29, 32].

Reconstruction of the division complexes in LLPS systems, both in bulk and encapsulated inside microfluidic microdroplets, has allowed analyzing the effect of microenvironments and confinement on the observed condensation under crowding conditions. SlmA and, more remarkably, the SlmA·SBS complexes share some of the preferred locations of FtsZ, asymmetric and largely dictated by its self-association state [32, 33], in the model LLPS systems mimicking intracellular microenvironments studied here. That is, they accumulate at the dextran or DNA rich phases, possibly entailing implications for their reactivity and for their differential recognition with other biomolecules and, most likely, strengthening their interaction with each other. Partition into the two mentioned phases is particularly pronounced in the case of the FtsZ·SlmA condensates, both in the absence and presence of SBS, which could be the result of a favored condensation related with an increase in the local concentration of their integrating elements in these phases.

Probably the most notable effect derived from liquid droplet formation on the location of the FtsZ·SlmA·SBS complexes is the observed tendency of the condensates to concentrate at or nearby the lipid boundary when encapsulated inside microdroplets. Although it was initially proposed that disassembly of FtsZ filaments by SlmA would occur in the nucleoid [34, 48], different models suggest the antagonist activity hampering FtsZ ring formation falls at the membrane [34, 36, 42], still near the nucleoid that is condensed occupying a large part of the cytoplasm [27]. It is intriguing, however, how a nucleoid-bound protein can counteract FtsZ ring formation at the membrane [34], and one of the hypothesis raised is that SlmA could be brought there through transertional linkages involving membrane encoding sequences near the SBS ones [42]. Our results suggest that the formation of crowding-driven FtsZ·SlmA·SBS condensates could also aid in membrane localization. There, SlmA may compete with the anchoring proteins FtsA and ZipA for their common target sequence, the C-terminal tail of FtsZ [36] (**Fig. 7**), and reinforce the function of the inhibitory Min System that operates through waves travelling across the membrane [49]. At the moment of division, chromosome segregation and migration towards the poles would decrease the amount of SlmA at midcell, that together with the accumulation of ZipA at this location [50] may favor competition driving FtsZ out of the inhibitory complexes [36, 51], thus allowing fibers formation and hence FtsZ ring assembly.

**Fig. 7.**
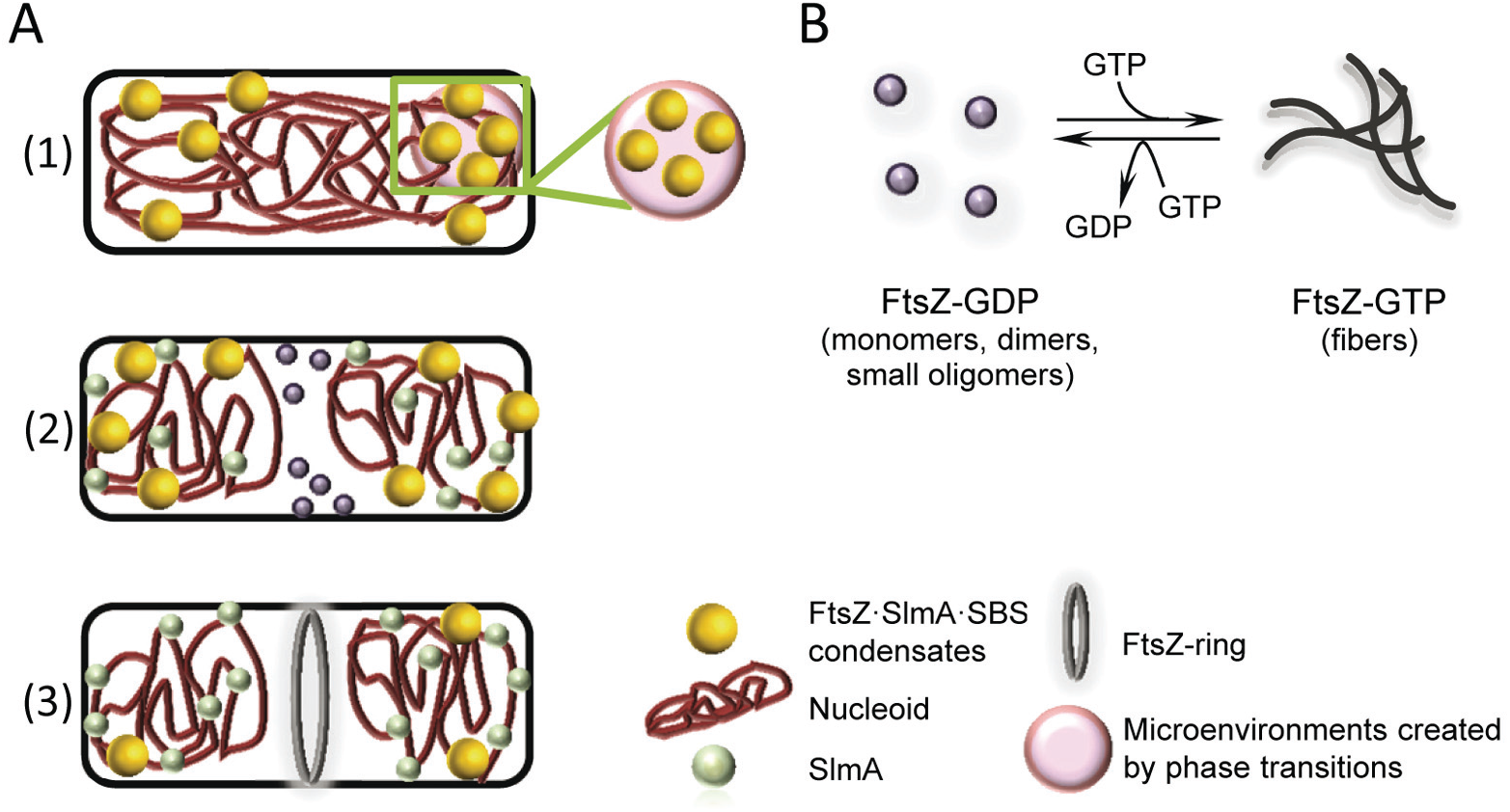
Scheme for the hypothetical influence of FtsZ·SlmA·SBS condensation on the action of SlmA over FtsZ in bacterial division. (**A**) (1) Under non division conditions, SlmA bound to its specific DNA target sequences (SBS) within the chromosome may recruit unassembled FtsZ, forming condensates in the crowded cytoplasm. The local concentration of the three elements in these condensates would be increased, likely enhancing their mutual recognition. Additionally, the nucleoprotein condensates may further accumulate in transient microenvironments resulting from crowding induced phase transitions in the cytoplasm. Condensation into liquid droplets may aid in the localization of the elements in the vicinity of the membrane, where SlmA would compete with the membrane anchoring proteins for FtsZ binding [36], and reinforce the inhibition of FtsZ fibers formation by other systems operating at the membrane [27]. If eventually FtsZ fibers are formed, they would be under the control of SlmA, which would limit their size and accelerate their disassembly to reorganize into FtsZ·SlmA·SBS condensates. (2) Under division conditions, chromosome segregation and migration towards the poles decrease the amount of SlmA at midcell [35] at the same time that the membrane anchor protein ZipA accumulates there [50], (3) likely competing for the interaction with FtsZ, that would leave the complexes with SlmA·SBS and, in the presence of GTP, form a membrane anchored FtsZ ring. In non-central regions, FtsZ would still be under the control of SlmA, which protects the chromosomes from scission by aberrant division ring formation. (**B**) Scheme of the selfassociation of FtsZ. In its GDP form, FtsZ is found as an ensemble of species of small size [45]. GTP binding induces its assembly into fibers that disassemble upon depletion of the nucleotide by FtsZ GTPase activity [26, 28].

The nucleoprotein condensates of FtsZ and SlmA are dynamic, allowing the incorporation of additional protein, the rapid evolution of the integrated FtsZ towards filaments in the presence of GTP, and its recruitment back into the liquid droplets upon GTP depletion. The dynamism and reversibility are hallmarks of liquid droplets and they appear to be particularly convenient for the role of SlmA as a spatiotemporal regulator of the formation of FtsZ filaments. Enhancement of the mutual interactions between the two proteins and the target SBS by their local accumulation within these condensates would ensure that, although SlmA·SBS does not completely block FtsZ assembly, the fibers eventually formed under its control are smaller and rapidly disassemble to condense again into FtsZ·SlmA·SBS liquid droplets. FtsZ within these condensates would remain active for assembly when and where required, and indeed super-resolution fluorescence imaging *in vivo* has provided evidences of FtsZ, upon Z-ring disassembly, persisting as patches that may act as precursors for its reassembly, also suggested to be involved in the formation of mobile complexes by recruitment of other binding partners [52]. Future efforts using these high resolution imaging technologies may aid to decipher the role of these transiently formed structures in division.

We propose that the modulation of FtsZ assembly by the specific nucleoprotein complexes of SlmA may be exerted through the generation of dynamic supramolecular condensates in which the protein can be accumulated in a non-filamentous manner, under the control of the antagonist. The tendency of these structures to localize at the lipid layers may be part of the mechanism by which SlmA exerts its antagonistic action at the membrane, and microenvironments transiently occurring in the cell may also modulate the behavior of the entire system by accumulating these liquid droplets in certain compartments changing their reactivity. The dynamic and reversible character of these condensates would endow plasticity to the spatiotemporal regulation of Z-ring formation by SlmA. Rather than to the few prokaryotic membraneless compartments so far identified, the condensates described here more closely resemble protein liquid droplets recently described in eukaryotes, opening the possibility of this kind of phase transition playing also a role in the regulation of bacterial processes. The reconstruction strategies in compartmentalized model cells applied here can greatly contribute to establish the possible consequences of these phase transitions in other systems as well.

## Methods

### Materials

GTP nucleotide, dextran 500, PEG 8 and other analytical grade chemicals were from Sigma. Polar extract phospholipids from *E. coli*, from Avanti Polar Lipids (Alabama, USA), were kept in chloroform at −20°C. Before use, a lipid film made by drying EcL in a Speed-Vac device or under a nitrogen stream was resuspended in mineral oil to the final concentration by two cycles of vortex plus 15 min sonication in a bath.

### Protein purification and labelling

FtsZ and SlmA were purified as described [35, 37, 45] and stored at −80°C until used. The proteins were covalently labelled in the amino groups with Alexa 488 or Alexa 647 carboxylic acid succinimidyl ester dyes (Molecular Probes/Invitrogen) as earlier stated [30, 37, 53], and stored at −80°C. The ratio of labelling of FtsZ and SlmA, ranging between 0.2-0.9 moles of fluorophore per mole of protein, was estimated from their molar absorption coefficients. For the experiments, protein solutions were equilibrated in 50 mM Tris-HCl, 300 mM KCl, 1 mM MgCl2, pH 7.5.

### Specific SBS oligonucleotides hybridization

Complementary single stranded oligonucleotides, either unlabeled or labeled with fluorescein in 5’ using the phosphoramidite chemistry, were purchased from IBA GmbH, and hybridized using a thermocycler as explained [37]. The fluorescein labelled oligonucleotide was hybridized with a 10% excess of the complementary unlabeled one. The double stranded oligonucleotides generated in this way contained the SBS consensus sequence specifically targeted by SlmA (SBS, 5’-AA**GTAAGTGAGCGCTCACTTAC**GT-3’, bases recognized by SlmA in bold) [35].

### Unspecific DNA fragmentation and purification

For its use as crowder in LLPS systems, salmon sperm DNA (Wako Pure Chemical Industries, Japan) was fragmented and purified as described [32], following a slightly modified phenol:chloroform:isoamyl alcohol extraction method [44]. The dried pellet was resuspended in 50 mM Tris-HCl, 300 mM KCl, pH 7.5. The DNA obtained, fragments of ≤ 300 bp [32], was kept at −20°C until used. DNA concentration was estimated from its dry weight after purification. Slight variations were found from batch to batch in the concentration at which phase separation with PEG was achieved, probably reflecting slight differences in DNA quantification, which did not affect the behavior of the division elements in this LLPS system.

### Preparation of phases for LLPS systems and labelling of PEG

Enriched phases were prepared by mixing and subsequent isolation of PEG 8 and dextran 500 or unspecific DNA in 50 mM Tris-HCl, 300 mM KCl, pH 7.5 at concentrations rendering phase separation, as described in detail [32, 33]. Fluorescent labelling of PEG was done as described [32].

### Preparation of bulk emulsions of PEG/dextran and PEG/DNA LLPS systems

The bulk emulsions were formed by thoroughly mixing PEG-rich and dextran 500-rich or PEG-rich and DNA-rich phases in a 3:1 volume ratio as described [32]. Proteins were directly added to this mixture and, when required, polymerization of FtsZ was triggered by diffusion of GTP directly added over the mixture of the two phases. Localization of proteins and of the double stranded SBS oligonucleotide in the LLPS system phases was evaluated from the colocalization of the labelled element with a tracer amount (1 μM) of PEG-Alexa 488 or PEG-Alexa 647 depending on the dye attached to the protein or SBS. Images were acquired with different combinations of dyes (FtsZ-, SlmA-Alexa 488 and SBS-Fl with PEG-Alexa 647, FtsZ-, SlmA-Alexa 647 with PEG-Alexa 488) with equivalent results (**Supplementary Figs. 5** and **6**). The concentrations of FtsZ, SlmA and SBS were 12, 5 and 1 μM, respectively, unless otherwise stated. The buffer for the aqueous solutions in all experiments was 50 mM Tris-HCl, 300 mM KCl, 1 mM MgCl2, pH 7.5.

### Diffusion of additional FtsZ into the preformed condensates with SlmA

Samples with FtsZ and SlmA ± SBS double stranded oligonucleotide containing 1 μM FtsZ labelled with Alexa 647 in the specified crowding agents were prepared and imaged before and after addition of 0.5-1 μM FtsZ-Alexa 488. The diffusion of FtsZ-Alexa 488 into the red labelled droplets was monitored with time. The concentrations of FtsZ, SlmA and SBS in these experiments were 12, 5 and 1 μM, respectively.

### Turbidity measurements

Turbidity of samples containing 12 μM FtsZ, 5 μM SlmA and 1 μM SBS in the presence and absence of Ficoll or dextran 500 (150 g/L) or PEG 8 (50 g/L) was determined at room temperature and 350 nm using a Varioskan Flash plate reader (Thermo). The absorbance of 200 μL solutions was measured every 10 minutes for 140 min and it was found to be stable during this time period. Reported values, average of 3 independent measurements ± SD, correspond to those recorded after 2 h incubation.

### Measurement of the partition of division elements in LLPS systems by fluorescence

Partition within the PEG/dextran mixture was calculated as described [32]. Briefly, tracer (0.5 μM Alexa 488 labelled proteins or fluorescein labelled SBS) and unlabeled species up to the concentrations specified (12, 5 and 1 μM FtsZ, SlmA and SBS, respectively) were gently added to the two phases in buffer in a 1:1 volume ratio. Mixture was allowed to phase separate and equilibrate for 30 min and, after centrifugation, phases were isolated and the fluorescence emission intensity of an aliquot of each phase measured in PolarStar Galaxy (BMG Labtech, GmbH, Germany) or Varioskan (Thermo) Plate Readers. Concentrations in the enriched phases were calculated by comparison with samples containing known amounts of tracer in the same phase. Control measurements proved tracer signals were in all cases linear with total concentration. Reported values correspond to the average of 3 independent measurements, 6 in the case of the samples with the three components, ± SD.

### Microfluidic chip fabrication

Chips were constructed by conventional soft lithographic techniques as earlier explained [33]. PDMS base SylgardTM 184 was mixed in a 10:1 (w/w) with curing agent (Dow Corning GmbH, Germany), degassed, decanted onto masters (design details described elsewhere [54]) and kept overnight at 65°C. Inlet and outlet holes were punched in the PDMS peeled from the master and channels sealed by a glass slide activating the surfaces by oxygen plasma (Diener electronic GmbH, Germany). For hydrophobic treatment of the chips, Aquapel (Pittsburgh Glass Works, LLC) was flushed in the channels and dried overnight at 65°C.

### Encapsulation of LLPS systems in microdroplets by microfluidics

Production of microdroplets by microfluidics was conducted basically as described [33]. Briefly, encapsulation was achieved by mixing a PEG 8 stream and another one with either dextran 500 or salmon sperm DNA in an approximately 1:1 volume ratio prior to the droplet formation junction. When stated, Alexa-647 or Alexa-488 labelled PEG (2 μM) was included in the PEG solution. FtsZ (12 or 25 μM) was added to one of the aqueous phases and SlmA (6 or 10 μM) with or without SBS (1-2 μM) was added to the other. When required, tracer amounts (2 μM) of the proteins labelled with the specified dye and of the SBS oligonucleotide labelled with fluorescein (2 μM) were added.

Comparable results were obtained with all dye combinations. When induction of FtsZ polymerization before encapsulation was required, the nucleotide GTP (2-4 mM) was included in the SlmA solution. The third stream supplied the mineral oil with the *E. coli* lipid mixture (20-25 g/L). In the particular case of the PEG/DNA LLPS system encapsulation, surfactant capability of the lipids seemed to be lower. Solutions were delivered at 120 μL/h (oil phase) and 5 and 7 μL/h (dextran and PEG aqueous phases, respectively), or 6 μL/h (both DNA and PEG aqueous phases), by automated syringe pumps (Cetoni GmbH, Germany). Droplets production in the microfluidic chip was monitored with an Axiovert 135 fluorescence microscope (Zeiss).

### Confocal microscopy measurements and data analysis

The microdroplets generated by microfluidics were visualized either on chip or after collection in silicone chambers (Molecular probes/Invitrogen) glued to coverslips. These chambers were also used to visualize the division complexes in the presence of crowding agents or in the LLPS systems. Images were obtained with a Leica TCS-SP2-AOBS or a Leica TCS-SP5-AOBS inverted confocal microscope with a HCX PL APO 63x oil immersion objective (N.A. = 1.4–1.6; Leica, Mannheim, Germany). Ar (488 nm) and He-Ne (633 nm) ion lasers were used to excite Alexa 488/Fluorescein and Alexa 647, respectively. Image J (National Institutes of Health, USA) was used to produce images and time-lapse movies, to measure the distribution of sizes and to obtain the intensity profiles of the liquid droplets applying the straight line tool of the software through their equatorial section.

## Acknowledgements

Authors thank W.T.S. Huck and A. Piruska (Radboud University, Nijmegen) for kindly providing the chips designs and silicon masters for microfluidics, N. Ropero for technical assistance in protein purification and labelling, M.T. Seisdedos and G. Elvira (Confocal Laser and Multidimensional Microscopy Facility, CIB-CSIC) for excellent support in imaging and the Technical Support Facility (CIB-CSIC) for invaluable input. This work was supported by the Fondo Europeo de Desarrollo Regional (FEDER) and the Agencia Estatal de Investigación (AEI); by the Spanish government (BFU2014-52070-C2-2-P and BFU2016-75471-C2-1-P, G. R.) and by the National Science Foundation (MCB-1715984, C. D. K.). M.L.-A. was supported by the European Social Fund (ESF 2014-2020).

## Author contributions

B.M., S.Z. and G.R. conceived the experimental work; B.M. and S.Z. analyzed results; B.M., S.Z., M.S.-S., M.R.-R. and M.L.-A. performed experimental work; B.M., S.Z., C.D.K. and G.R. discussed the results and wrote the manuscript.

## Additional information

**Supplementary Information** accompanies this paper.

